# DLK inhibition has sex-specific effects on neuroprotection and locomotor recovery after spinal cord injury

**DOI:** 10.1101/2025.10.17.683054

**Authors:** John C. Aldrich, Samantha M. Alman, Sydney E. Lee, Ashley R. Scheinfeld, Chelsea C. Zhang, Averi L. Pike, Fiona C. Bremner, Olivia Calderon, Sunil Goodwani, William J. Ray, Andrew D. Gaudet

**Author notes:** Corresponding Author: Andrew Gaudet, Department of Psychology University of Texas at Austin, 108 E. Dean Keeton Street, Stop A8000, Seay Building, Room 4.208 Austin, TX 78712, USA.

## Abstract

Spinal cord injury (SCI) causes devastating functional deficits, in part due to neuroinflammation, oxidative stress, and excitotoxicity that drive death of lesion-adjacent viable neurons. Dual leucine zipper kinase (DLK) is a neuron-enriched kinase that responds to cellular stress by activating the c-Jun N-terminal kinase (JNK) pathway, driving both stress-responsive gene expression and neuronal apoptosis. We hypothesized that SCI would robustly activate DLK signaling and that acute pharmacological inhibition of DLK would suppress JNK pathway activation, thereby enhancing neuroprotection and locomotor recovery in our mouse model of moderate contusion SCI. Using western blotting, we observed that SCI induced strong and sustained activation of the JNK pathway in the injured spinal cord starting at 4 hours post-injury through 7 days. Complementary analysis of single-nucleus RNA-seq revealed that DLK expression is highly enriched in neurons across all injury phases. Following SCI, neurons exhibited robust, time-dependent upregulation of multiple DLK-responsive transcripts, consistent with sustained pathway activation during the acute and subacute periods. Systemic treatment with the selective DLK inhibitor IACS-52825 effectively suppressed intraspinal JUN activation in a dose-dependent manner. However, unexpectedly, treatment delayed functional recovery and expanded lesion volume by 71% in male mice with no significant effect in females. These findings highlight the complex roles of DLK signaling after SCI, revealing a need to understand the sex-specific molecular mechanisms that modulate injury outcomes. Future studies should further optimize timing, location, and cellular targeting of DLK therapeutic strategies to improve neuroprotection and neurologic recovery after SCI.

## INTRODUCTION

Spinal cord injury (SCI) causes loss of motor, sensory, and autonomic function, creating chronic locomotor deficits and neuropathic pain that can permanently burden affected individuals (Ahuja et al., 2017; Anjum et al., 2020). Following the initial mechanical insult, inflammation and excitotoxicity create a stressful microenvironment within the injured spinal cord that leads to the loss of otherwise viable neurons. This secondary damage limits the long-term potential for functional recovery (Gaudet & Fonken, 2018; Lukacova et al., 2021; Pang et al., 2022). Preventing secondary damage and neuronal loss during the early stages of injury remains a central goal of research involving rodent models of SCI, yet effective therapies that improve long-term outcomes remain elusive. As such, there is growing interest in targeting neuron-intrinsic pathways that regulate stress responses and determine cell survival after injury.

Dual leucine zipper kinase (DLK, encoded by *Map3k12*) is a critical neuron-intrinsic sensor of cellular stress, rapidly activated in response to axonal injury, metabolic stress, and extracellular signals such as inflammation and excitotoxicity (Pozniak et al., 2013; Shin et al., 2019; Tedeschi & Bradke, 2013; Watkins et al., 2013). Upon activation, DLK signals through the JNK (Jun N-terminal kinase) pathway, leading to phosphorylation of the transcription factor JUN and induction of stress-responsive transcriptional programs. The functional outcomes of DLK activation are highly context dependent, driving neuronal apoptosis and degeneration in some settings while promoting axonal regeneration in others (Jin & Zheng, 2019; Tedeschi & Bradke, 2013). In the adult central nervous system—where regenerative capacity is inherently limited—the consequences of modulating DLK signaling after injury have not been fully explored. Combined neuronal deletion of DLK and its close paralog LZK (Leucine zipper-bearing kinase, *Map3k13*) promotes neuroprotection in a traumatic brain injury model (Welsbie et al., 2019) while suppressing *Pten*-dependent axonal regeneration after hemisection SCI (Saikia et al., 2022). Thus, these kinases have dual and context-dependent roles after CNS injury. Notably, astrocyte-specific LZK deletion increases lesion size and impairs functional recovery after SCI (Chen et al., 2018; Hemati-Gourabi et al., 2025), demonstrating a non-neuronal role for this protein family in regulating injury outcomes. Although pharmacological inhibition of DLK has not been evaluated after SCI, DLK-targeting drugs improve neuronal survival and limit degeneration in models of glaucoma, amyotrophic lateral sclerosis, and Alzheimer’s disease, highlighting the potential of this neuroprotective strategy following SCI (Le Pichon et al., 2017; Patel et al., 2015; Siu et al., 2018; Welsbie et al., 2013; Wlaschin et al., 2018).

Given DLK’s central role in neuronal stress signaling, selective small molecule inhibitors targeting this kinase have been developed to modulate JNK-mediated neuronal stress signaling (Bu et al., 2023; Oetjen & Lemcke, 2016; Patel et al., 2015). IACS-52825, a highly selective, brain-penetrant DLK inhibitor has demonstrated robust *in vivo* efficacy and therapeutic potential in models of peripheral neuropathy (Le et al., 2023; Ma et al., 2021). In this study, we evaluated the effects of IACS-52825 treatment in a mouse model of moderate contusion SCI. We hypothesized that SCI would induce DLK signaling in the injured spinal cord and that acute pharmacological inhibition would enhance neuroprotection and promote locomotor recovery. Western blotting and single-nucleus RNA-Seq analyses showed that JUN and other targets of DLK signaling are strongly activated after SCI and remain elevated through the acute to subacute phase. IACS-52825 suppressed JUN activation in a dose-dependent manner. However, contrary to our hypothesis, treatment delayed locomotor recovery and increased lesion size in male mice, with no effect in females, highlighting previously unrecognized sex-specific consequences of targeting this pathway.

## METHODS

### Animals and housing

All housing, surgery, and postoperative care procedures were approved by The University of Texas at Austin Institutional Animal Care and Use Committee (Protocol AUP-2021-00171). Adult (12-16 weeks old) male and female C57BL/6J mice (Jackson Laboratory, Stock 000664) were housed in sex-matched pairs, provided standard chow and filtered tap water *ad libitum*, and maintained on a 12:12 light/dark cycle. Mice in all treatment groups were numbered randomly to ensure experimenters were blind to group assignments.

Sample sizes for each experiment are as follows: western blot time course: n = 3 males and 3 females (sham, 4 hr), n = 2 males and 3 females (1 dpo, 7 dpo). IACS-52825 dose-response western blot: sham/vehicle n = 3 males and 3 females, SCI/vehicle n = 4 males and 4 females, SCI/3 mg/kg n = 4 males and 4 females, SCI/10 mg/kg n = 5 males and 4 females, SCI/30 mg/kg n = 5 males and 4 females. 48 hr IACS-52825 i.p. western blot: n = 4 males and 4 females per treatment group. 28 dpo locomotor recovery: vehicle n = 5 males and 6 females, IACS-52825 n = 5 males and 6 females. Lesion analysis: vehicle n = 4 males and 5 females, IACS-52825 n = 4 males and 5 females.

### Surgeries and postoperative care

All surgeries were performed as previously described (Gaudet et al., 2016; Lee et al., 2023; Scheff et al., 2003) between Zeitgeber time (ZT) 2–6. Mice were anesthetized with isoflurane inhalation anesthesia (1.5%; MWI Animal Health, Cat. 502017) and received prophylactic buprenorphine hydrochloride (0.075 mg/kg; MWI Animal Health, Cat. 060969) immediately prior to surgery. A dorsal T9 laminectomy was performed, with the periosteum but not the dura removed. Sham mice received no further injury. For SCI mice, a moderate contusion injury (65 kdyn, 0 s dwell) was delivered at T9 using the Infinite Horizon impactor (Precision Systems and Instrumentation). Incisions were closed using sutures and wound clips. Postoperative care included daily subcutaneous injections of Ringer’s solution (2, 2, 1, 1, 1 mL on postoperative days 1–5) to prevent dehydration and manual bladder expression twice daily, which was reduced to once daily once bladder function returned. Mice were monitored at least once daily for signs of infection or abnormal recovery.

### Drug formulation and delivery

IACS-52825 was provided by the M.D. Anderson Belfer Neurodegeneration Consortium and is described previously (Le et al., 2023; Ma et al., 2021). For initial dose-response characterization, IACS-52825 was formulated fresh in 0.5% (w/v) methylcellulose (FujiFilm Wako, Cat. 133-17815) (10 mL/kg) and administered once by oral gavage 3 hours post-surgery at 3, 10, or 30 mg/kg. The compound was suspended in vehicle, vortexed, sonicated in a water bath (10 min) and homogenized using a BioSpec Tissue-Tearor (5–7 min, half speed). Lower doses were prepared by diluting the 30 mg/kg formulation in 0.5% methylcellulose. Solutions were kept at room temperature with continuous mixing until administration. Vehicle control mice received 0.5% methylcellulose alone.

For continual dosing, IACS-52825 was prepared in a vehicle of polyethylene glycol 400 (PEG 400; MilliporeSigma, Cat. 95904) and polysorbate 80 (Tween 80; MilliporeSigma, Cat. P4780) in sterile water (40:10:50, v/v/v; 10 mL/kg) and administered intraperitoneally at 30 mg/kg beginning 3 h post-SCI and then at 2, 4, and 6 dpo (ZT2). The compound was dissolved in the PEG 400/Tween 80 mixture by vortexing (30 s, maximum speed) and sonicating in a water bath sonicator at room temperature (10–30 min) until fully dissolved. Sterile water was then added, and the solution was vortexed vigorously for 30–60 s before dosing. Vehicle control mice received PEG-400/Tween 80/sterile water alone.

### Locomotor testing

Hindlimb motor function was assessed using the Basso Mouse Scale (BMS; Basso et al., 2006) prior to surgery and at 1, 4, 7, 10, 14, 21, and 28 days post-injury (dpo). Recovery was scored on the BMS scale of 0 (complete paralysis) to 9 (normal motor function) by two observers blinded to treatment conditions. For statistical analysis (see below) left and right hindlimb scores were included as a fixed effect; since there was no significant difference between sides (F_1,297_ = 0.23; p = 0.632), the average of both was used for visualization purposes.

### Western blotting

Western blotting was performed as previously described (Goodwani et al., 2020; Le et al., 2023; Ma et al., 2021). Briefly, mice were euthanized with Euthasol (200–270 mg/kg; Virbac AH, Inc, Cat. 011355), transcardially perfused with ice-cold PBS, and a 5 mm spinal cord segment centered at the T9 lesion epicenter was dissected and flash frozen on dry ice. Tissue was lysed in radioimmunoprecipitation assay buffer (RIPA; 150 mM NaCl, 1.0% IGEPAL CA-630, 0.5% sodium deoxycholate, 0.1% SDS, 50 mM Tris, pH 8.0) containing protease and phosphatase inhibitors (ThermoFisher, Cat. PI78440), homogenized on ice, and centrifuged at 14,000 × *g* for 30 min at 4 °C. Protein concentrations were determined by DC Protein Assay (Bio-Rad, Cat. 5000111), diluted in loading buffer (Bio-Rad, Cat. 1610737) with 5% 2-mercaptoethanol (Bio-Rad, Cat. 1610710), denatured by heating to 95°C for 10 min, 25 µg was loaded and separated by SDS-PAGE on 4-12% Bis-Tris gels (NuPAGE, ThermoFisher, Cat. NP0321) using 1× MES running buffer (ThermoFisher, Cat. NP0002) and transferred to nitrocellulose membranes via wet transfer in NuPAGE Transfer Buffer (ThermoFisher, Cat. NP0006-1) with 10% methanol. Membranes were blocked with Odyssey Blocking Buffer (LI-COR Biosciences, Cat. 927-40000) and incubated overnight at 4°C with primary antibodies diluted in blocking buffer against phospho-JUN (pJUN; Abcam, Cat. ab32385; 1:100), JUN (Cell Signaling Technology, Cat. 9165; 1:250), phospho-MAP2K4 (Cell Signaling Technologies, Cat. 9156; 1:100), and GAPDH (MilliporeSigma, Cat. MAB374; 1:5000). Membranes were washed three times with TBS-T (Tris-buffered saline with 0.1% Tween 20) between incubation steps and incubated with IRDye-conjugated secondary antibodies (LI-COR Biosciences, Cat. 926-68021 and 926-68020; 1:5000) for 1 h at room temperature. Immunoreactive bands were visualized using an Odyssey CLx Imaging System (LI-COR Biosciences) and quantified with Image Studio software.

### Tissue collection and processing

Mice were euthanized via an intraperitoneal injection of Euthasol (200–270 mg/kg; Virbac AH, Inc.) followed by transcardial perfusion with cold 1× phosphate-buffered saline (PBS; pH 7.4) and 4% formaldehyde in PBS. Spinal cords (5 mm segments including the injury epicenter and 2.5 mm rostral and caudal) were dissected and post-fixed overnight in cold 4% formadehyde (MilliporeSigma, Cat. P6148) before transferring to 30% sucrose the following day. Tissue was embedded in optimal cutting temperature compound (OCT; Fisherbrand, Cat. 23-730-571), flash frozen on dry ice, and sectioned at 12 µm on a cryostat. Serial cross sections were collected onto Superfrost Plus slides (Fisherbrand, Cat. 12-550-15) and stored at −20°C until use. Mice were randomized so that tissue collection, handling, and subsequent analysis were performed in a blinded manner.

### Immunohistochemistry, imaging, and lesion analysis

Immunohistochemistry was performed as previously described (Aldrich et al., 2024; Gaudet et al., 2016). Briefly, slides were brought to room temperature, encircled with a hydrophobic barrier pen, washed in PBS, and blocked for 1 h at room temperature in 10% normal donkey serum (NDS; Jackson, Cat. NC9624464) in PBS with 0.2% Triton X-100 (PBTx). Sections were incubated overnight at 4 °C with Rabbit anti-GFAP (1:500; Cell Signaling Technology, Cat. 3670) diluted in 10% NDS in PBTx. The following day, slides were washed, incubated for 2 h at room temperature with Donkey anti-Rabbit Alexa Fluor 488 (1:500; ThermoFisher, Cat. A21206) and DAPI (1:10,000), washed again, mounted in Immu-Mount (Epredia, Cat. 9990402), cover-slipped, and sealed.

Spinal cord sections were imaged on a Nikon Ni-E widefield epifluorescence microscope using the “acquire large image” function in Nikon Elements (AR 5.21.02) at the Center for Biomedical Research Support Microscopy and Flow Cytometry Facility at UT Austin (RRID:SCR_021756). Nine overlapping 10× fields were stitched to form each composite image. For each mouse, the epicenter section—defined as the section with the largest GFAP-negative lesion area—plus five rostral and five caudal sections (every 20th section, spaced 240 µm apart) were imaged and analyzed. Lesion area and total cross-sectional area were quantified in ImageJ (v2.1.0; Fiji distribution) based on GFAP staining (Schindelin et al., 2012; Schneider et al., 2012).

### Single-nucleus RNA sequencing analysis

For snRNA-Seq analyses, we used data from the *Tabulae Paralytica* atlas (Gene Expression Omnibus accession GSE234774; Skinnider et al., 2024). This large-scale resource comprises single-nucleus transcriptomic profiles from multiple injury models, severities, timepoints, ages, sexes, and spinal cord regions. For our analyses, we specifically used data from the lesion epicenter of adult mice (8-15 weeks old) following moderate mid-thoracic (T10) crush spinal cord injury. We analyzed female samples (n = 3 biological replicates per timepoint except 7 dpo, n = 2) at 1, 4, 7, 14, 30, or 60 days post-injury, plus uninjured controls. For sex comparisons, we also analyzed male samples at 7 dpo (n = 3).

In the original study, the spinal cord injury site was dissected immediately following euthanasia and frozen on dry ice. Nuclei were isolated by mechanical homogenization in sucrose buffer using a dounce homogenizer followed by density gradient centrifugation to obtain purified nuclei. Libraries were prepared using the 10X Genomics Chromium Single Cell Gene Expression Kit v3.1 targeting 10,000 nuclei per sample and sequenced to approximately 75,000 reads per nucleus. After quality control and ambient RNA removal, the authors performed batch correction using Harmony followed by iterative Leiden clustering in Seurat to identify cell types and subtypes based on canonical marker gene expression.

We downloaded processed count matrices, feature tables, and metadata from GEO and assembled into a Seurat object using the *ReadMtx*, *read.delim*, and *CreateSeuratObject* functions in the *Seurat R* package (v5.3.0; Hao et al., 2024). To quantify expression of *Map3k12* (DLK) and *Map3k13* (LZK), we calculated the percentage of cells with ≥ 1 count for each gene within each layer (layer1) for all biological replicates, excluding categories with fewer than 25 cells per replicate. We also fit binomial generalized linear mixed models (GLMMs) using *lme4* (v1.1-37; Bates, Mächler, Bolker, & Walker, 2015) and *broom.mixed* (v0.2.9.6) for each layer and for all subtypes (layers 1–5), including a log10-transformed total count term to control for library size and replicate ID to account for between-sample variation. Counts were aggregated within each layer using *Seurat’s AggregateExpression* function, and differential gene expression was tested with *DESeq2* (v1.42.1; Love, Huber, & Anders, 2014). For the time course, we used a likelihood ratio test (LRT) to identify genes whose expression varied across time points, and a Wald test to compare each time point to the uninjured condition for neurons only. For the 7 dpo analysis, we used a Wald test to compare gene expression between males and females for each layer. Uncorrected p-values from *DESeq2* were collected across all layers and cell types, and a global false discovery rate (FDR) adjustment was applied to obtain adjusted p-values (Benjamini & Hochberg, 1995).

To identify coordinated expression patterns among DLK-responsive genes, we performed hierarchical clustering on row-scaled (z-score normalized) log₂ fold change values using Euclidean distance and complete linkage, partitioning genes into four clusters using *ComplexHeatmap* (v2.18.0; Gu, Eils, & Schlesner, 2016). For each cluster, we calculated eigengenes—summary profiles representing the dominant pattern of coordinated expression—via principal component analysis (PCA) on the z-scored expression matrix, similar to approaches used in weighted gene co-expression network analysis (Langfelder & Horvath, 2008). The first principal component (PC1) was extracted and oriented to match the mean cluster trajectory, then scaled to unit variance to facilitate visualization across clusters.

### Statistics, data analysis, and visualization

For western blot time course data, two-way ANOVAs (type III sum of squares) were performed for each target using the *Anova* function from the *car* package (v3.1-3; Fox & Weisberg, 2019), with group (sham, 4 h, 1 d, 7 d post-injury), sex, and their interaction as fixed effects, and blot number included as a covariate to control for variation across different blots. *Post-hoc* comparisons were performed using estimated marginal means via the *emmeans* library (v1.11.1; Lenth, 2024) comparing each SCI timepoint to sham controls and adjusting p-values for multiple testing via the FDR method. For dose–response results, type III ANOVA was again used with dose (as a numeric variable), sex, and their interaction as fixed effects along with blot number as a covariate. *Post-hoc* contrasts were assessed using *emmeans* to compare each dose to the vehicle control.

For analysis of repeated measures data (BMS scores across multiple days and lesion size at serial tissue sections), we fit linear mixed-effects models using *lmerTest* (v3.1-3; Kuznetsova, Brockhoff, & Christensen, 2017), including mouse ID as a random effect. Upon detecting a significant *treatment x sex x dpo* interaction in the BMS data, BMS scores and lesion size were analyzed separately for males and females, with fixed effects including treatment, time (or section order), and their interactions; for BMS scores, foot was also included as a fixed effect. When significant interactions were detected, post-hoc contrasts were performed with *emmeans* using FDR correction. For lesion volume, we performed two-tailed Welch’s t-tests to compare treatment groups within sex using the *t.test* function from the *stats* package (v4.3.2).

All data analysis and visualization were performed in *R* (v4.3.2; R Core Team, 2022) using *RStudio* (v2025.5.1.513; RStudio Team, 2020). Data wrangling was done primarily with *tidyverse* (v2.0.0), especially *dplyr* (v1.1.4). Figures were generated with *ggplot2* (v3.5.2) except for heatmaps, which were created with *ComplexHeatmap* (v2.18.0l; Gu, Eils, & Schlesner, 2016). Multi-panel figures were assembled using *cowplot* (v1.1.3), *ggpubr* (v0.6.0), *legendry* (v0.2.2), and *patchwork* (v1.3.0). Color palettes were optimized for color-blind accessibility with *scico* (v1.5.0) (Crameri et al., 2024). Additional tools included *ggrepel* (v0.9.6) for labeling volcano plots*, ggbeeswarm* (v0.7.2) for jitter plots, and *ggtext* (v0.1.2) for formatted text. Data import/export used *readr* (v2.1.5), *readxl* (v1.4.5), and *openxlsx* (v4.2.8). The *ragg* package (v1.4.0) was used for high-quality graphics rendering, and *conflicted* (v1.2.0) was used to manage function conflicts between packages.

## RESULTS

### SCI activates the DLK/JNK pathway in the lesion epicenter

DLK is a central mediator of neuronal stress signaling and has emerged as a promising therapeutic target in models of axonal injury and neurodegeneration. DLK activates downstream responses via the MAP2K4,7–JNK–JUN signaling axis (**Fig. 1A**), leading to transcriptional activation of stress response and apoptotic pathways (Patel et al., 2015; Tedeschi & Bradke, 2013). To assess pathway activation in our spinal cord injury (SCI) model, we performed moderate (65 kdyn) T9 contusion injuries in adult male and female C57BL/6 mice and collected epicenter tissue at 4 hours, 1 day, and 7 days post-injury. Western blotting was used to quantify phosphorylation of MAP2K4 and JUN, as well as total JUN expression levels, which increases following JNK pathway activation (**Fig. 1B–D**; **Fig. S1**).

**Figure 1:**
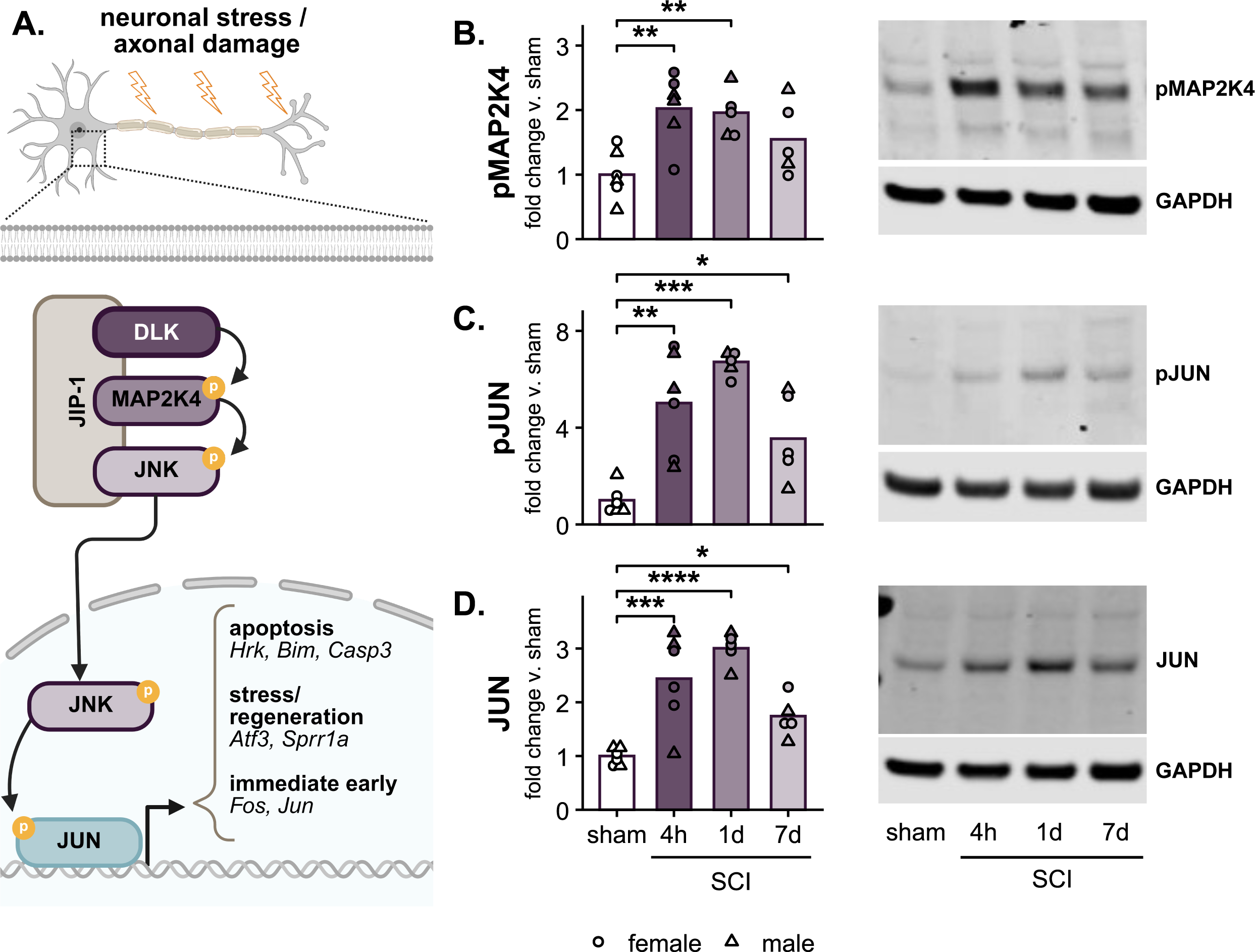
SCI activates DLK/JNK signaling in the lesion epicenter. **(A)** Schematic of the DLK signaling cascade. Extracellular stress activates DLK, which phosphorylates MAP2K4, leading to phosphorylation of JNK. JNK translocates to the nucleus and activates the transcription factor JUN. **(B-D)** Western blot quantification and representative images of phospho-MAP2K4 (pMAP2K4) **(B),** phospho-JUN (pJUN) **(C)**, and total JUN **(D)** protein levels in the lesion epicenter at 4 hours, 24 hours, and 7 days post-SCI, compared to sham. GAPDH was used as a loading control, and values are expressed relative to sham. Each point represents an individual mouse (n = 2-3/sex/group): females (circles), males (triangles). Asterisks indicate significant post-hoc comparisons to sham following a significant group effect in two-way ANOVA (**Table S1** & **S2**): *p < 0.05, **p < 0.01, ***p < 0.001, ****p < 0.0001. Full blot images are provided in **Fig. S1**. Pathway diagram created in BioRender (2025) https://BioRender.com/r0daq7w.

Phosphorylated MAP2K4 (pMAP2K4) was elevated by 4 hours post-injury and peaked at ∼2-fold above sham at 1 dpo (p = 0.007; **Fig. 1B**). Phospho-JUN increased similarly by 4 hours and peaked at ∼6.8-fold above sham at 1 day (p = 0.0003), with elevated levels persisting through 7 days (p = 0.028; **Fig. 1C**). Total JUN protein also rose by 4 hours, peaked at 3-fold at 1 day, and remained elevated at 7 days (p = 1.5 x 10^-5^ and p = 0.016; **Fig. 1D**). There were no detectable main effects of sex (F_1,13_ ≤ 1.2, p ≥ 0.29) or sex × group interactions (F_1,13_ ≤ 0.21, p ≥ 0.89) for targets examined (**Table S1** & **S2**). Overall, these data show sustained activation of DLK/JNK signaling during the first week following SCI.

### DLK expression is enriched in neurons across SCI timepoints

To evaluate cell-type specificity of DLK expression in the spinal cord, we re-analyzed single-nucleus RNA-sequencing (snRNA-Seq) data from the *Tabulae Paralytica* atlas, which captured approximately 10,000 nuclei per mouse from the lesion epicenter at multiple timepoints following SCI, as well as uninjured controls (Skinnider et al., 2024). In this approach, nuclei from 2-3 mice per condition were isolated, sequenced, and grouped into clusters based on similarity in gene expression patterns. Clusters were then assigned cell type identities—ranging from broad classes like neurons and astrocytes to fine-grained subtypes such as *Rreb1^+^ Zim1^+^* dorsal excitatory neurons—based on the expression of established marker genes. This dataset allows us to analyze gene expression at the single-cell level or within aggregated groups of cells (i.e., pseudobulk analysis). Expression of DLK (i.e., *Map3k12*) was generally low and sparse across all populations. To simplify interpretation and minimize variability across replicates, we grouped cells into four time bins—uninjured, acute (1–4 dpo), subacute (7–14 dpo), and chronic (30–60 dpo)—and calculated the proportion of cells with detectable *Map3k12* expression (>0 counts) within each major cell class. Neurons consistently showed the highest proportion of DLK-expressing cells across all phases (**Fig. 2A**).

**Figure 2:**
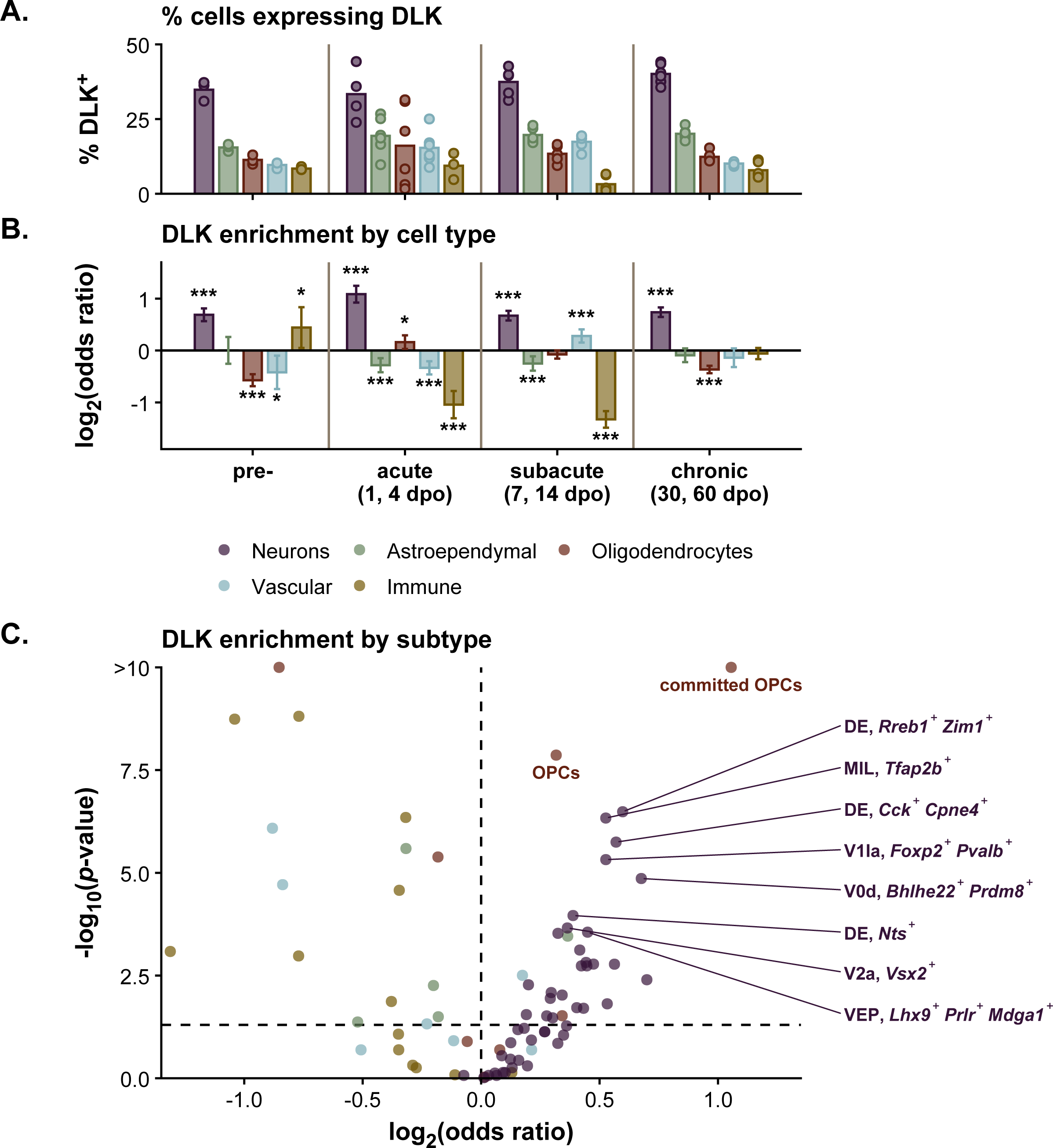
DLK expression is enriched in spinal neurons before and after SCI. **(A)** Percentage of DLK-expressing cells (*Map3k12* > 0) across major cell types and SCI phases, derived from published single-nucleus RNA-Seq data. Each point represents the average per biological replicate; n = 2-3 female mice per time. Timepoints are grouped into uninjured, acute (1–4 days post-injury), subacute (7–14 days), and chronic (30–60 days). **(B)** Cell type–specific enrichment of DLK expression, estimated using a binomial generalized linear model. Plotted are log₂ odds ratios (OR) with 95% confidence intervals. Asterisks indicate statistically significant enrichment compared to all other cell types (*p < 0.05, **p < 0.01, ***p < 0.001, ****p < 0.0001). **(C)** Subtype-level enrichment of DLK expression using the same modeling approach, aggregated across timepoints. Each point represents a distinct cell subtype; x-axis shows -log₁₀(FDR-adjusted *p*-value), y-axis shows log₂ odds ratio. The ten most significantly enriched subtypes (positive log₂ OR) are labeled: DE (dorsal excitatory), MIL (medial excitatory, lateral), V1 (ventral inhibitory interneurons), V2 (ventral excitatory interneurons), V0 (ventral commissural interneurons), VEP (ventral excitatory projection neurons), OPC (oligodendrocyte precursor cells).

To formally test whether DLK expression is preferentially associated with neurons, we used a binomial generalized linear model to estimate the likelihood of *Map3k12* detection within each major class relative to all others. Neurons were the only population with consistently elevated odds of DLK expression across all injury phases (odds ratio ≥ 1.6, FDR-adjusted p ≤ 2 x 10^-27^). In contrast, other classes—including oligodendrocytes, immune cells, and astroependymal populations—generally exhibited depletion of DLK, with only transient enrichment observed at isolated timepoints (**Fig. 2B, Table S3**). We note that this analysis does not preclude DLK expression in other cell types but rather indicates that neurons show the highest likelihood of expressing detectable *Map3k12* relative to other spinal cord populations.

Next, we applied the same modeling approach to all annotated cell subtypes in the dataset, including granular distinctions among neuronal, immune, vascular, and glial populations (**Fig. 2C, Table S4**). Subtypes were pooled across timepoints to reduce dimensionality and increase the representation of rarer cell populations. Of the 47 neuronal subtypes examined, 26 showed significantly elevated likelihood of DLK expression (odds ratio ≥ 1.14, FDR-adjusted p ≤ 0.034), and none were significantly depleted. In contrast, only five non-neuronal subtypes showed significant enrichment, including two oligodendrocyte progenitor populations (odds ratio ≥ 1.27, FDR-adjusted p ≤ 0.030). A complementary pseudobulk RNA-Seq analysis using a similar one-versus-all approach mirrored these results, with *Map3k12* significantly upregulated in 73% of neuronal subtypes along with the same two oligodendrocyte populations (**Fig. S2**, **Table S5**). RNA *in situ* hybridization from the Allen Brain Institute (Allen Institute for Brain Science, 2008; Lein et al., 2007) revealed *Map3k12* mRNA enrichment specifically in spinal cord grey matter cells (**Fig. S2**), consistent with neuronal enrichment. Together, these findings indicate that DLK is preferentially expressed in neurons, supporting its relevance as a neuron-intrinsic target for modulating stress responses and promoting neuroprotection following SCI.

DLK and LZK are closely related paralogs sharing 70% sequence homology with nearly identical catalytic domains (Sakuma et al., 1997; Welsbie et al., 2017). Even highly selective kinase inhibitors would likely exhibit some degree of cross-reactivity with LZK due to this structural similarity. We therefore examined LZK expression patterns to characterize the cellular distribution of this related kinase. Similar to DLK, LZK (*Map3k13*) showed enrichment in neurons across injury phases, with some expression also observed in oligodendrocyte lineage cells (**Fig. S2, S3**).

### SCI induces expression of DLK-responsive genes

To assess DLK pathway activation in neurons after SCI, we performed pseudobulk differential expression analysis using aggregated snRNA-Seq data from all spinal cord neurons. Counts were summed per replicate and timepoint, and gene-level changes relative to uninjured controls were quantified using *DESeq2* (Love et al., 2014). We evaluated a curated set of genes whose transcriptional expression is regulated by DLK activity, including canonical DLK/JNK pathway downstream targets and genes previously identified as DLK-dependent in optic nerve crush injury using DLK knockout mice (Watkins et al., 2013). These “DLK-responsive genes” represent transcripts whose injury-induced expression was abolished or attenuated in the absence of DLK.

Of the 381 DLK-associated transcripts detected in the neuronal pseudobulk dataset, 58.3% showed significant changes over time (FDR-adjusted p < 0.05 based on likelihood ratio testing; **Table S6, Fig. 3A**). These findings are consistent with our western blot results from bulk spinal cord tissue (**Fig. 1B–D**) and indicate that neurons mount a robust, time-dependent transcriptional response involving established DLK/JNK targets after SCI.

**Figure 3:**
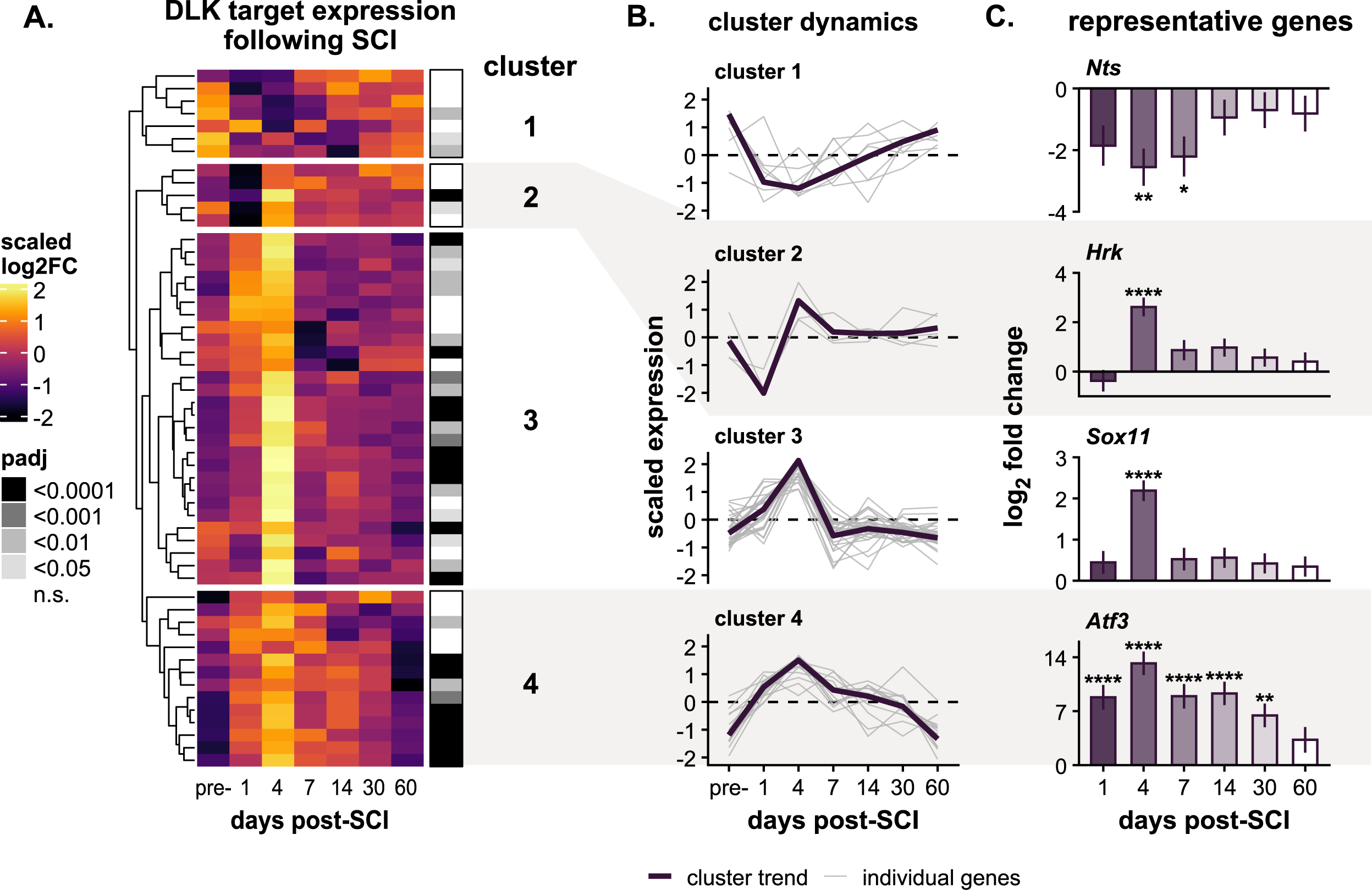
DLK-responsive genes are transcriptionally induced in neurons after SCI. **(A)** Pseudobulk differential expression analysis of DLK-associated genes in spinal cord neurons across timepoints following SCI. Rows represent individual genes and columns represent discrete timepoints from uninjured through 60 days post-injury. Color reflects row-scaled (z-score) log₂ fold change values calculated from pairwise comparisons to uninjured controls. Genes are grouped by hierarchical clustering using Euclidean distance and complete linkage. Cluster identities are shown numerically to the right, and FDR-adjusted p-values from likelihood ratio testing are indicated by grayscale. **(B)** Temporal expression dynamics for each of the four gene clusters. Individual gene trajectories (grey) and cluster eigengenes (purple) are shown as scaled expression values across timepoints. Eigengenes represent the first principal component of expression patterns within each cluster, capturing the dominant coordinated trend. **(C)** Expression profiles of representative DLK-responsive genes from each cluster showing log₂ fold change (mean ± standard error) relative to uninjured controls at each timepoint. Asterisks indicate significant differences compared to uninjured (*p < 0.05, **p < 0.01, ***p < 0.001, ****p < 0.0001; FDR-adjusted).

Expression patterns among these DLK-associated transcripts grouped into four clusters. To visualize the dominant temporal dynamics within each cluster, we calculated cluster eigengenes—summary profiles that capture the primary pattern of coordinated expression change across all genes in each cluster (**Fig. 3B**). Representative individual genes from each cluster are shown in **Fig. 3C**. Cluster 1 genes, such as *Nts*—involved in synaptic signaling—were higher at baseline and were suppressed post-injury, with varying degrees of return toward pre-injury levels by 30–60 days. Cluster 2 genes showed a brief spike at 4 days post-injury after early suppression; this included *Hrk*, a pro-apoptotic factor, which increased ∼2.6-fold at 4 dpo (p = 2 x 10^-9^). Cluster 3, the largest group, consisted of transcripts that peaked at 4 dpo and returned to baseline by 7 days. These included well-known regeneration-associated genes *Sox11*, *Gadd45a*, and *Ecel1*. Cluster 4 genes—including *Atf3*, *Clic4,* Ddit3 (i.e., CHOP, an ER stress response gene)—rose at 1 dpo, peaked at 4 dpo, and remained elevated through at least 30 days. *Atf3* was among the most strongly induced, with >13-fold upregulation at 4 dpo (p = 3.2 x 10^-15^).

Together, these findings demonstrate that SCI causes neurons to upregulate multiple DLK-responsive transcripts, with coordinated transcriptional activation evident throughout the acute post-injury phase (1–7 dpo), consistent with DLK/JNK pathway engagement in neurons following injury.

### IACS-52825 treatment suppresses JUN activation following SCI

Given the sustained activation of DLK/JNK signaling observed in neurons during the acute phase of SCI, we next tested whether inhibition of this pathway could suppress downstream JUN activation *in vivo*. In a dose-response study, adult mice received oral IACS-52825 at doses that achieve low, moderate, and high CNS target engagement (3, 10, or 30 mg/kg) or vehicle control 3 hours after moderate T9 contusion SCI. Spinal cord tissue was collected 24 hours later for western blot analysis of phospho-JUN and total JUN.

IACS-52825 treatment reduced SCI epicenter phospho-JUN levels in a dose-dependent manner (F_3,25_ = 10.7, p = 6.3 x 10^-4^; **Fig. 4A, Fig. S4, Table S7**). All three IACS-52825 doses reduced epicenter pJUN relative to vehicle, with the 30 mg/kg group showing the largest effect (effect size = −6.5; p = 1 x 10^-4^, **Table S8**). Similar results were observed for total JUN (**Fig. 4B**), with 30 mg/kg IACS-52825 most potently decreasing epicenter JUN (ANOVA F_3,25_ = 15.7, p = 7.8 x 10^-6^; *post hoc* effect size = - 2.9; p = 8.5 x 10^-6^).

**Figure 4:**
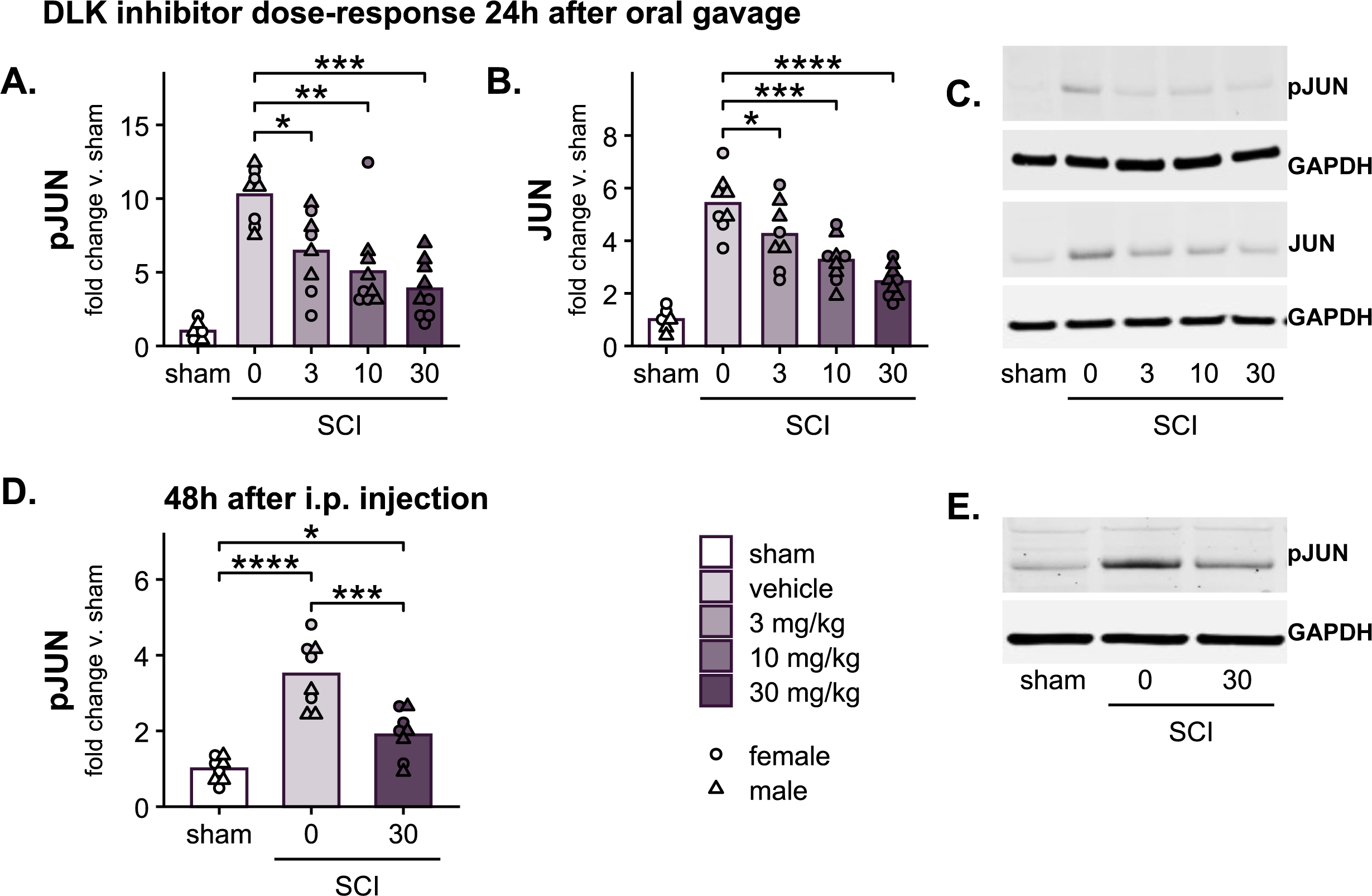
IACS-52825 treatment suppresses JUN phosphorylation after SCI. **(A)** Phospho-JUN (pJUN) levels in spinal cord epicenter tissue 24 hours after SCI and oral administration of IACS-52825 (3, 10, or 30 mg/kg) or vehicle. IACS-52825 treatment produced a dose-dependent reduction in pJUN compared to vehicle. **(B)** Total JUN levels from the same samples, showing a similar dose-dependent decrease. **(C)** Representative western blot images for the dose-response experiment. **(D)** pJUN levels 48 hours after SCI and a single intraperitoneal (i.p.) injection of IACS-52825 (30 mg/kg) or vehicle. **(E)** Representative western blot images for the i.p. experiment. All values are normalized to GAPDH and expressed relative to sham. Circles represent females, triangles represent males (3–4 mice per sex per group). Asterisks indicate significant post-hoc comparisons (*p < 0.05, **p < 0.01, ***p < 0.001, ****p < 0.0001) following a significant group effect in two-way ANOVA. Full blot images are provided in **Fig. S4**.

To facilitate repeated dosing across the full acute phase (1–7 dpo), we reformulated IACS-52825 for intraperitoneal (i.p.) injection and adopted an every-other-day schedule. This route was selected to minimize stress associated with repeated oral gavage (Balcombe et al., 2004; Larcombe et al., 2019; Vanhecke et al., 2024). To test efficacy, mice received a single 30 mg/kg i.p. dose 3 hours post-injury, and tissue was collected 48 hours later. As with oral delivery, i.p. IACS-52825 reduced epicenter pJUN levels relative to vehicle (ANOVA F_2,18_ = 32, p = 1.2 x 10^-6^; *post hoc* p = 1.1 x 10^-4^; **Fig. 4D, Fig. S4, Table S9** & **S10**). These results suggest that systemic IACS-52825 administration is an effective means of suppressing JNK pathway activation after SCI.

### IACS-52825 delays locomotor recovery in male mice after SCI

To assess the functional consequences of DLK/JNK inhibition, mice received IACS-52825 (30 mg/kg) or vehicle starting 3 hours post-SCI, with subsequent injections every 48 hours through 6 days post-injury (**Fig. 5A**). Locomotor function was evaluated using the Basso Mouse Scale (BMS) at 1, 4, 7, 10, 14, 21, and 28 dpo.

**Figure 5:**
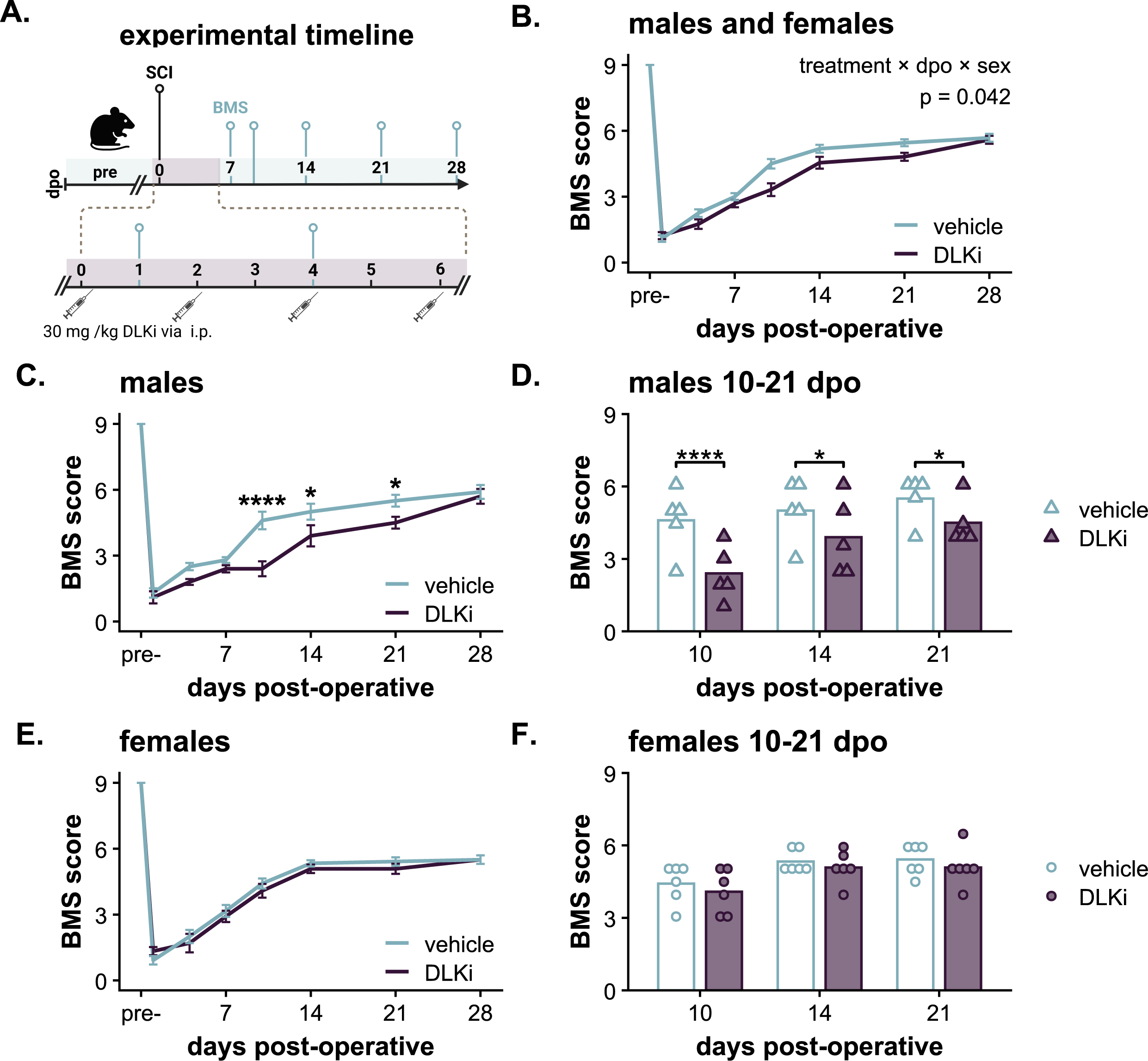
IACS-52825 treatment after SCI impairs locomotor recovery in males but has no significant effect in females. **(A)** Experimental timeline. Mice received IACS-52825 (DLKi; 30 mg/kg) or vehicle beginning 3 hours post-injury, with repeat dosing every 48 hours through 6 days post-SCI. BMS scoring was performed at the indicated timepoints, and tissue was collected at 28 days post-injury. **(B)** DLKi treatment does not improve BMS scores when analyzing males and females combined (vehicle vs. DLKi; females: *n* = 6 per group, males: *n* = 5 per group). Values reflect mean of left and right hindlimb scores ± SEM. A significant treatment × day × sex interaction was detected via linear mixed-effects model. **(C)** In males, IACS-52825 treatment led to significantly lower BMS scores at 10–21 days post-injury. **(D)** Individual male scores during this interval. **(E)** DLKi treatment had no significant effect on BMS scores in female mice overall or at specific timepoints **(F)**. Asterisks indicate significant *post-hoc* comparisons between treatment groups (FDR-adjusted). (*p < 0.05, **p < 0.01, ***p < 0.001, ****p < 0.0001). Timeline Created in BioRender. (2025) https://BioRender.com/9528xxc.

IACS-52825 treatment selectively impaired locomotor recovery in male mice during the early post-injury period. Longitudinal analysis of BMS scores revealed a significant treatment × day × sex interaction (F_7,297_ = 2.1, p = 0.042; **Fig. 5B, Table S11**). When stratified by sex, IACS-52825-treated males exhibited significantly lower BMS scores at 10, 14, and 21 dpo, with the largest effect observed at 10 dpo (estimate = - 2.2, p = 4.7 x 10^-6^; **Fig. 5C, Table S12**). At this timepoint, vehicle-treated males averaged 4.6 ± 0.4—consistent with the onset of occasional plantar stepping—while drug-treated males averaged 2.4 ± 0.3, indicating a delay in recovery. By 10 dpo, 4 of 5 vehicle-treated males exhibited plantar stepping, compared to only 1 of 5 in the IACS-52825 group. Despite these early differences, males from both treatment groups reached similar endpoints by 28 dpo, averaging a score of ∼6, which corresponds to frequent to consistent plantar stepping with at least some forelimb–hindlimb coordination (**Fig. 5D**). In contrast, female mice showed no significant difference in recovery between treatment groups at any timepoint (**Fig. 5E, F**). When comparing vehicle-treated animals, there were no significant differences in locomotor recovery between males and females (main effect of sex: F_1,9_ = 0.16, p = 0.7; sex × day interaction: F_7,148_ = 1.38, p = 0.22), suggesting similar baseline injury responses across sexes. Overall, these results indicate that systemic IACS-52825 treatment does not enhance locomotor recovery after SCI, and instead transiently delays locomotor recovery in male mice.

### IACS-52825 treatment increases lesion size in male mice after SCI

To evaluate the histological impact of IACS-52825 treatment, we analyzed 12 μm cryosections spanning the lesion epicenter at 240 µm intervals using fluorescence immunohistochemistry. Lesion and spared tissue areas were quantified using widefield fluorescence microscopy of GFAP-immunolabeled sections (**Fig. 6A**). The lesion was defined as the central GFAP-negative region, and spared tissue was calculated as the remaining GFAP-positive area expressed as a percentage of the total cross-sectional area.

**Figure 6:**
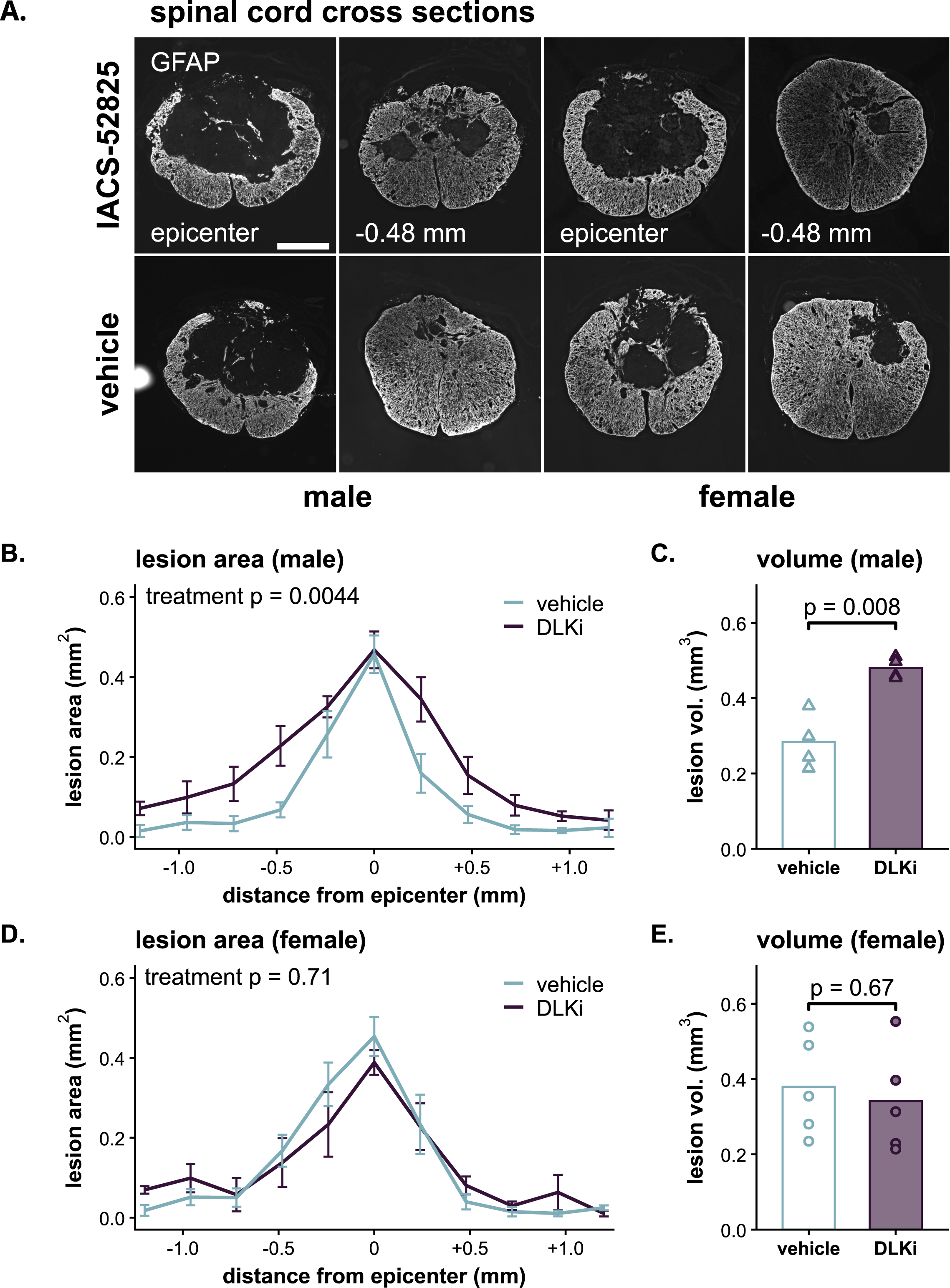
Systemic IACS-52825 delivery has sex-dependent effects on neuroprotection after SCI. **(A)** Representative GFAP-immunolabeled spinal cord cross sections 28 days after dorsal midline contusion SCI from vehicle- and IACS-52825-treated male and female mice. Images from the epicenter and 480 µm rostral are shown. Lesion area was defined as the central GFAP-negative region, and spared tissue was calculated as the remaining GFAP-positive area. Scale bar indicates 0.4 mm. (**B**) IACS-52825 (DLKi) increased lesion area in male sections spanning ±1.2 mm from the epicenter—11 sections per animal spaced 240 µm apart. (**C**) Lesion volume, estimated by interpolating area measurements, was greater in treated males. (**D**) DLKi had no significant effect on lesion area in females. (**E**) Lesion volume in females was similar between groups. females: *n* = 5 per group, males: *n* = 4 per group. P-values in panels **B** and **D** reflect main effects of treatment from linear mixed-effects models (treatment × position), while those in **C** and **E** are from unpaired t-tests comparing total lesion volume. Error bars represent SEM. See **Figure S5** for spared tissue quantification.

Consistent with our BMS results, IACS-52825 treatment exacerbates tissue damage in male mice following SCI. When analyzing both sexes (**Fig. S5A, Table S13**), we observed a treatment × sex interaction for both lesion area (F_1,14_ = 5.7, p = 0.03) and percent spared tissue (F_1,14_ = 5.3, p = 0.04), prompting follow-up analysis stratified by sex. In male mice, IACS-52825 treatment increased lesion area relative to vehicle-treated controls (**Fig. 6B**; LMM main effect of treatment: F_1,6.5_ = 18.3, p = 0.004). Lesion volumes interpolated from these measurements showed a 0.2 mm³ increase in the IACS-52825 treatment group—equivalent to a 71% increase relative to vehicle (unpaired t-test: t_3.8_ = 5.04, p = 0.008; **Fig. 6C**). Quantification of spared tissue mirrored these results (F_1,6_ = 14, p = 0.009; **Fig. S5E, F**).

No differences between vehicle or IACS-52825 treatment were observed in female SCI mice for lesion area (F_1,8_ = 0.11, p = 0.75; **Fig. 6D, Table S13**), lesion volume (t_8_ = −0.45, p = 0.67; **Fig. 6E**), or spared tissue (F_1,8_ < 0.01, p = 0.9; **Fig. S5G, H**). Furthermore, vehicle-treated males and females showed similar lesion areas (main effect of sex: F_1,7_ = 1.52, p = 0.26; sex × position interaction: F_10,57_ = 0.6, p = 0.81) and percent spared tissue (main effect of sex: F_1,7_ = 0.18, p = 0.68; sex × position interaction: F_10,57_ = 0.47, p = 0.90), indicating comparable baseline histological outcomes between sexes following injury. These findings align with the locomotor recovery data, indicating that systemic IACS-52825 treatment fails to promote tissue preservation after SCI and may worsen outcomes in males. Overall, the data show that while IACS-52825 effectively targets the JNK pathway, it does not confer neuroprotection or functional benefit following spinal cord injury.

## DISCUSSION

DLK is an evolutionarily conserved neuronal stress sensor studied across various model systems (e.g., *Drosophila*, *C. elegans*, and mammals) in diverse injury and disease contexts (Le Pichon et al., 2017; Tedeschi & Bradke, 2013; Xiong et al., 2010; Yan et al., 2009). Despite its therapeutic potential, DLK remains relatively uncharacterized in the adult spinal cord, and its activation dynamics after SCI have not been well defined. DLK regulates neuronal stress and apoptosis primarily through the JNK pathway (Ghosh et al., 2011; Miller et al., 2009). As such, we found that contusion SCI induces robust phosphorylation of the JNK targets MAP2K4 and JUN in the epicenter throughout the acute-to-subacute post-injury phase. snRNA-Seq analysis suggests DLK mRNA enrichment in spinal cord neurons and showed that SCI induces downstream transcriptional targets over a similar timeframe. We hypothesized that acute inhibition of the DLK/JNK pathway via IACS-52825 would enhance neuroprotection and improve locomotor recovery after SCI. Although IACS-52825 effectively reduced post-SCI JUN activation, it failed to improve functional outcomes and instead led to worse recovery in male mice. These results demonstrate that IACS-52825 can effectively modulate JNK signaling after SCI, but that systemic treatment has sex-specific effects and is unlikely to support functional recovery.

Phosphorylation of DLK targets MAP2K4 and JUN increases rapidly by 4 hours post-SCI and remains elevated through 7 days (**Fig. 1**), mirroring DLK pathway activation in the optic nerve following both direct and indirect injury (Huntwork-Rodriguez et al., 2013; Watkins et al., 2013; Welsbie et al., 2019; Wong & Benowitz, 2022). Unlike DLK (**Fig. 2**), JUN is broadly expressed across neuronal and non-neuronal populations and can be activated by several upstream kinases. Nevertheless, studies using global DLK knockout in optic nerve injury models show that the characteristic injury-induced phosphorylation of JUN is almost completely lost without DLK, indicating that in CNS trauma contexts JUN activation is largely dependent on DLK signaling (Le Pichon et al., 2017; Watkins et al., 2013). In the spinal cord, DLK is highly enriched in neurons, and pharmacological inhibition with IACS-52825 markedly reduces pJUN after SCI (**Fig. 4**), suggesting that DLK signaling drives SCI-induced JNK pathway activation.

DLK activation triggers a transcriptional program that coordinates neuronal stress responses, apoptosis, and regenerative growth, largely via JUN-dependent regulation of downstream targets (Bu et al., 2023; Tedeschi & Bradke, 2013). In the ONC model, Watkins et al. identified a set of DLK-dependent genes whose injury-induced expression was abolished or attenuated in DLK knockout mice (Watkins et al., 2013). Using a previously published snRNA-Seq dataset, we found that many of these same genes—including *Atf3, Sox11*, and the pro-apoptotic factors *Hrk and Ddit3* (aka *CHOP*)—are induced in neurons after SCI (**Fig. 3**). Additionally, several canonical DLK pathway components absent from the Watkins list (e.g., *Jun* and other AP-1 transcription factors *Fos*, and *Fosl1*) were also induced in SCI epicenter. Together, these findings indicate that SCI engages a DLK-dependent transcriptional network closely paralleling that seen in the optic nerve, encompassing both stress-adaptive and pro-apoptotic elements.

In many models of neuronal injury and neurodegeneration, DLK loss-of-function or pharmacological inhibition confers neuroprotection, including paradigms such as optic nerve injury, excitotoxicity, and tauopathy (Le Pichon et al., 2017; Pozniak et al., 2013; Welsbie et al., 2013). DLK also plays a critical role in neural development, regulating axon growth, neuronal migration, programmed apoptosis, and axon degeneration during circuit refinement and maturation (Ghosh et al., 2011; Hammarlund et al., 2009; Miller et al., 2009; Tedeschi & Bradke, 2013; Yan et al., 2009). Other work has shown that DLK/JNK signaling is a key regulator of axonal regeneration, with injury-induced stress responses triggering transcriptional programs that can drive both axon growth and degeneration depending on context (Itoh et al., 2009; Jin & Zheng, 2019; Shin et al., 2019). This dual capacity suggests that the impact of inhibiting these pathways likely varies with injury type, timing, and the relative contribution of protective versus regenerative responses.

Direct investigation of DLK in SCI remains limited. To date, the only *in vivo* study found that neuronal DLK/LZK double knockout reduced corticospinal tract axon regeneration in a *Pten*^⁻^*^/^*^⁻^ background and suppressed spontaneous sprouting in wild-type mice, 18 weeks after dorsal hemisection SCI (Saikia et al., 2022). These effects were absent when either kinase was deleted alone, suggesting functional redundancy in the ability of DLK and LZK to enhance axon plasticity. Based on this and broader evidence that DLK can drive both degeneration and regeneration, we hypothesized that acute-to-subacute inhibition—when the post-SCI environment is most stressful—would suppress early maladaptive stress responses while sparing later regenerative processes. In our model, treatment with the IACS-52825 during this early window reduced JUN phosphorylation (**Fig. 4**) but did not improve locomotor recovery (**Fig. 5**) or neuroprotection (**Fig. 6**).

This unexpected outcome may reflect opposing effects of DLK inhibition—limiting apoptosis but also constraining stress-adaptive and repair pathways—that offset each other’s impact on functional recovery even at the acute-to-subacute timepoint. Our transcriptomic analysis supports this possibility, with important caveats. The transcripts we identified as DLK-responsive were defined based on altered expression in DLK knockout models (Watkins et al., 2013) and represent genes regulated by DLK activity rather than necessarily direct DLK/JNK targets. Moreover, stress-responsive and regeneration-associated transcriptional networks involve multiple converging pathways, of which DLK/JNK is just a prominent example (He et al., 2022; Zheng & Tuszynski, 2023). With these qualifications, SCI induces both pro-apoptotic (e.g., *Hrk, Ddit3*) and stress-adaptive (e.g., *Atf3, Sox11*) programs in spinal neurons (**Fig. 3**), suggesting that DLK activation engages parallel programs with potentially opposing functional consequences. Although axon plasticity is minimal in the injured adult spinal cord, many regeneration-associated factors are also linked to stress adaptation and may contribute to neuronal survival or repair (Mahar & Cavalli, 2018). Acute IACS-52825 treatment reduced JUN phosphorylation but may have simultaneously suppressed these protective responses, contributing to the absence of functional improvement.

Importantly, our findings should not be taken to imply that all DLK inhibition is inherently detrimental. Rather, they highlight the limitations of broad kinase inhibition in this context. Recent work has shown that disrupting DLK’s axonal localization can reduce pro-degenerative signaling while preserving beneficial transcriptional outputs (Zhang et al., 2025), suggesting that more targeted strategies may better dissociate these dual aspects of DLK function.

DLK and LZK are closely related paralogs sharing ∼70% overall sequence identity, with nearly identical catalytic domains (86% identity; Sakuma et al., 1997; Welsbie et al., 2017). As such, any small molecule kinase inhibitor targeting DLK would be likely exhibit some cross-reactivity with LZK. IACS-52825 demonstrates binding Kd values of 1.3 nM for DLK and 59 nM for LZK, with minimal engagement of over 400 other kinases tested in a comprehensive selectivity screen (Le et al., 2023). This indicates greater than 45-fold selectivity for DLK over LZK. However, given the CNS exposures achieved at the 30 mg/kg dose used in this study and the dose-response relationship observed in our experiments (**Fig. 4**), meaningful LZK engagement alongside DLK inhibition is likely. To the extent that DLK and LZK serve overlapping roles in neuronal stress signaling, this cross-reactivity is not necessarily problematic. Indeed, inhibiting both kinases would be expected to further suppress JNK pathway activation, which was the intended pharmacological effect. However, DLK and LZK also have distinct and non-canonical roles (Adula et al., 2022; Saikia et al., 2022; Welsbie et al., 2013, 2017). Furthermore, recent work demonstrates that astrocyte-specific LZK deletion increases lesion size and impairs functional recovery after SCI (Chen et al., 2018; Hemati-Gourabi et al., 2025). If IACS-52825 inhibits LZK in astrocytes, this could counteract any neuroprotective benefit in neurons or even contribute directly to the detrimental effects observed in male mice.

Beyond failing to provide neuroprotective benefit, IACS-52825 treatment actively worsened outcomes in male mice—delaying locomotor recovery and expanding lesion size (**Fig. 5, 6**). Although our western blot analyses were not specifically designed to assess sex differences (e.g., only 3–4 animals per sex per group were used), they revealed no obvious differences in DLK/JNK activation post-SCI or in sensitivity to IACS-52825 (**Fig. 1, 4**). This suggests that any sex differences in JNK signaling are either modest or obscured by cellular heterogeneity. Given the limited attention to sex as a variable in SCI research (Lee et al., 2023; Stewart et al., 2020), few datasets directly compare male and female transcriptional responses. The *Tabulae Paralytica* single-cell atlas (Skinnider et al., 2024) is a notable exception, but still only includes a single male–female comparison at 7 dpo. Overall, the authors report very few sex differences in cell-type proportions, transcriptional programs, or neurological outcomes, noting that sex accounted for the least variance among all comparisons. Upon revisiting this dataset to specifically evaluate DLK/JNK pathway components, we unexpectedly found that DLK (i.e., *Map3k12*) is significantly upregulated in male microglia relative to females (log₂ fold change = 5.7, p = 0.006; **Fig. S6, Table S14**). This was the only significant sex difference in pathway expression detected across cell types. This raises the possibility that non-neuronal DLK contributes to differential sensitivity between sexes. As discussed above, astrocyte-specific deletion of LZK causes lesion expansion and impaired locomotor recovery after SCI (Chen et al., 2018; Hemati-Gourabi et al., 2025), providing precedent for DLK-family kinases influencing recovery through non-neuronal mechanisms. More definitive insights into neuronal and non-neuronal DLK function after SCI will require further studies using temporally and cell type–specific genetic approaches. The unexpected, male-specific effects observed here suggest that sex may critically shape the outcomes of DLK-targeted interventions and should be considered an essential variable in any future mechanistic and translational work.

## CONCLUSION

Our study established that the selective DLK inhibitor IACS-52825 reduced phosphorylation of targets including JUN in SCI lesion epicenter and that DLK is enriched in neurons within the spinal cord. Despite the promising spatiotemporal distribution of DLK and the effectiveness of the inhibitor, systemic delivery of IACS-52825 had no significant benefit on neuroprotection or locomotor recovery—systemic IACS-52825 worsened neuroprotection and delayed locomotor recovery in males, but not females. Overall, the anatomical and functional effects of IACS-52825 were minimal or negative, which contradicts our original hypothesis. Future studies should extend these findings by exploring the spatiotemporal effects of DLK inhibition—e.g., by refining timing, cellular specificity, or lesion-relative location of DLK inhibition. Another effective approach may involve targeting signaling mediators upstream or downstream of DLK, which may provide more precise modulation of specific cells or unidirectional tuning of a crucial process (e.g., cell survival or axon plasticity).

Our results highlight that the DLK pathway has an important role in regulating recovery after SCI, and in a sex-specific manner. Future research will help crystallize our understanding of the role of the DLK pathway after CNS injury. Ultimately, unravelling the intricacies of DLK signaling could lead to new neuroprotective therapies that improve recovery after SCI.

## ACKNOWLEDGEMENTS

We thank the Animal Resources Center (ARC) husbandry staff at the Health Discovery Building for excellent animal care as well as the Center for Biomedical Research Support Microscopy and Flow Cytometry Facility at UT Austin (RRID:SCR_021756).

## FUNDING

Support for this work was provided by the Belfer Neurodegeneration Consortium as well as the National Institute of Neurological Disorders And Stroke of the National Institutes of Health under Award Number R01NS131806 (ADG). The content is solely the responsibility of the authors and does not necessarily represent the official views of the National Institutes of Health.

## DISCLOSURES

The authors have no competing interest to disclose.

## TRANSPARENCY, RIGOR, & REPRODUCIBILITY SUMMARY

All animal procedures were approved by The University of Texas at Austin IACUC (AUP-2021-00171). Mice were randomly assigned to treatment groups at the time of surgery, and behavioral scoring, tissue collection, and histological analyses were performed by investigators blinded to treatment and, when possible, to sex.

All experimental procedures—including spinal cord injury, drug preparation and administration, and postoperative care—followed established protocols as detailed in the Methods. Statistical analyses were conducted in R (v4.3.2) using linear mixed-effects models or ANOVA where appropriate. Model assumptions were checked through visual inspection and other standard diagnostics to confirm validity.

Single-nucleus RNA-seq data were obtained from the Tabulae Paralytica dataset (GEO accession GSE234774). No custom algorithms or hardware were used. Complete statistical outputs are provided in the Supplemental Materials. Code is available upon request, and data will be made publicly available upon publication at OSC-SCI.org.

## AUTHORSHIP CONTRIBUTION STATEMENT

**JCA:** Conceptualization; Data curation; Formal analysis; Investigation; Methodology; Project administration; Visualization; Writing – original draft; Writing – review & editing. **SMA:** Writing – original draft; Writing – review & editing. **SEL:** Investigation; Project administration. **ARS:** Investigation. **CCZ:** Investigation; Writing – original draft. **ALP:** Investigation. **FCB:** Investigation. **OC:** Investigation. **SG:** Investigation; Formal analysis; Supervision. **WJR:** Conceptualization; Funding acquisition; Supervision; Writing – review & editing. **ADG:** Conceptualization; Funding acquisition; Investigation; Supervision; Writing – original draft; Writing – review & editing.

## Notes

### Competing Interest Statement

The authors have declared no competing interest.

### Summary of Updates

Figures 1, 4, and 6 have been updated with representative images. The introduction, results, and discussion have been revised to expand on the potential for and implications of IACS-52825 cross-reactivity with LZK, a close paralog of DLK. Analysis of LZK expression was added to Figures S2 and S3.

