## Supplemental Figures for "DLK inhibition has sex-specific effects on neuroprotection and locomotor recovery after spinal cord injury"

### Supplementary Figures S1-S6

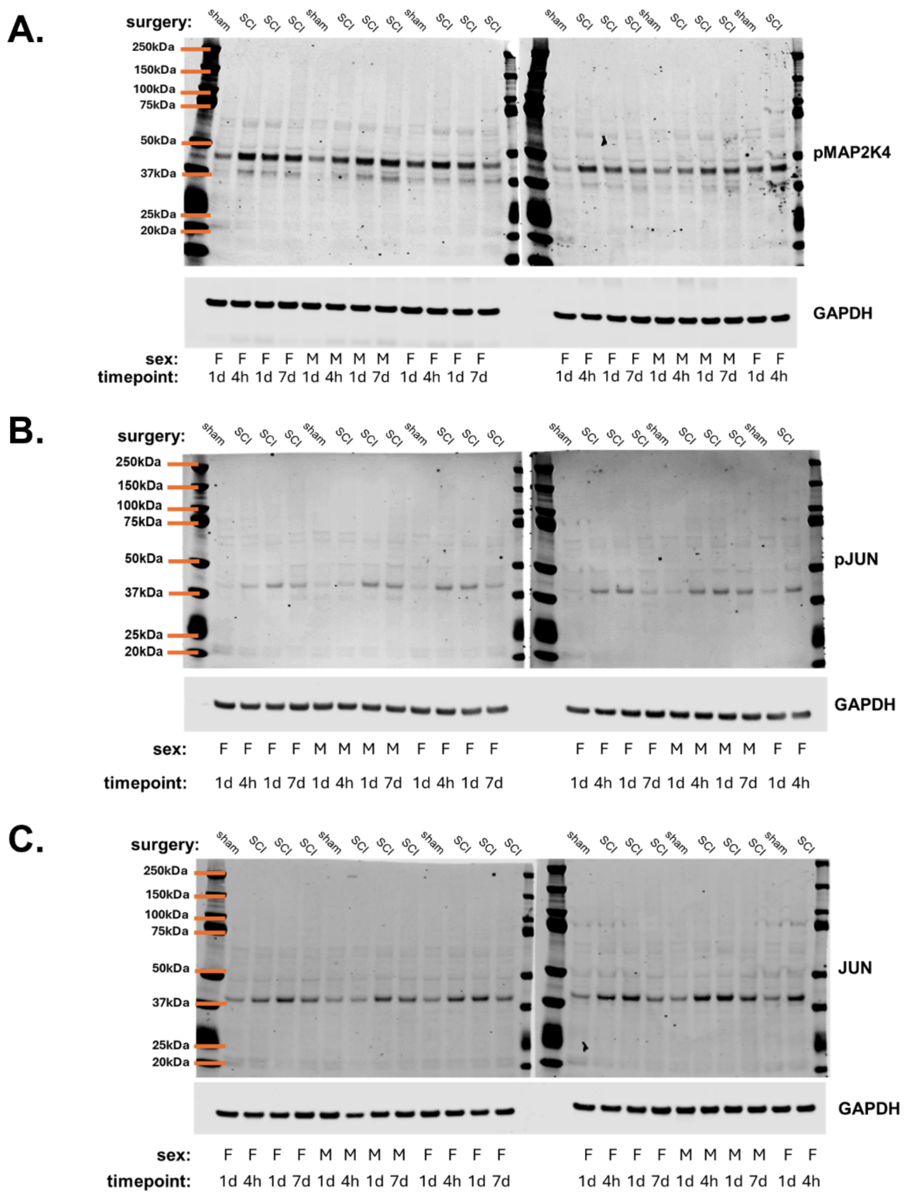


**Figure S1: Western blot images for DLK/JNK pathway proteins after SCI.**
Representative immunoblots for phospho-MAP2K4 (**A**), phospho-JUN (**B**), and total JUN (**C**) from spinal cord lesion epicenter tissue collected at 4 hours, 1 day, and 7 days post-SCI or sham surgery. GAPDH was used as a loading control for each blot. Molecular weight markers are indicated in kilodaltons (kDa). Above each blot are annotations for surgical group (SCI or sham); below each lane are annotations for sex (F = female, M = male) and timepoint (4h, 1d, 7d). Quantification of normalized and scaled band intensities is shown in **Fig. 1**.

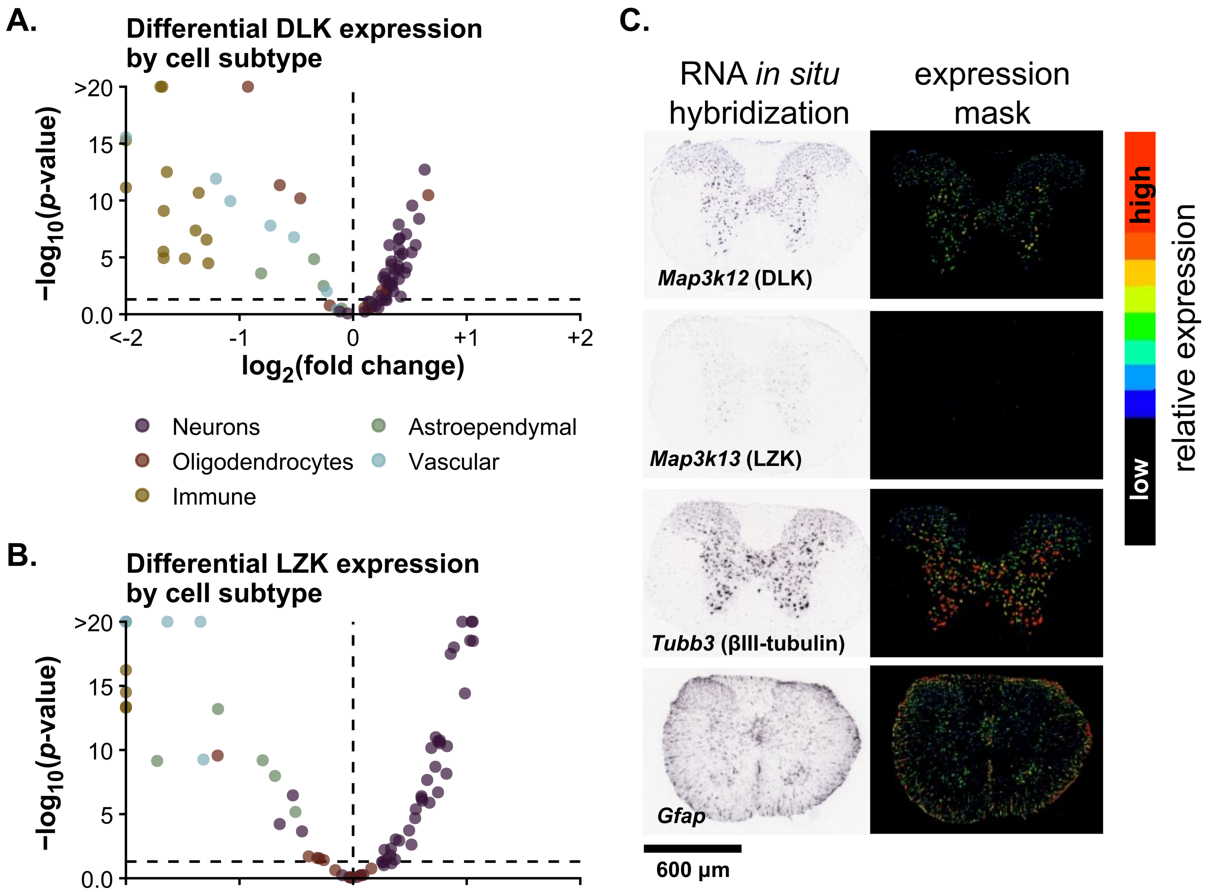


**Figure S2: DLK expression is enriched in neurons across cell subtypes. (A, B)** Volcano plots showing subtype-level enrichment of DLK (Map3k12; A) and LZK (Map3k13; **B**) expression using a one-versus-all differential expression approach applied to pseudobulk RNA-Seq data. Each point represents a distinct cell subtype; the x-axis shows -$\log_{10}$(FDR-adjusted p-value), and the y-axis shows $\log_2$ fold change of expression relative to all other subtypes. The ten subtypes with the most significant enrichment are labeled. **(C)** In situ hybridization images from the Allen Spinal Cord Atlas showing expression patterns of *Map3k12*, *Map3k13*, *Tubb3* (neuronal marker), and *Gfap* (astrocyte marker) in adult mouse spinal cord. Left panels show chromogenic *in situ* hybridization; right panels show expression masks indicating signal intensity from low (black) to high (red). Allen Spinal Cord Atlas, mousespinal.brain-map.org;
*Map3k12*: mousespinal.brain-map.org/imageseries/show.html?id=100012869
*Map3k13*: mousespinal.brain-map.org/imageseries/show.html?id=100024501
*Tubb3*: mousespinal.brain-map.org/imageseries/show.html?id=100013057
*Gfap*: mousespinal.brain-map.org/imageseries/show.html?id=100036944.

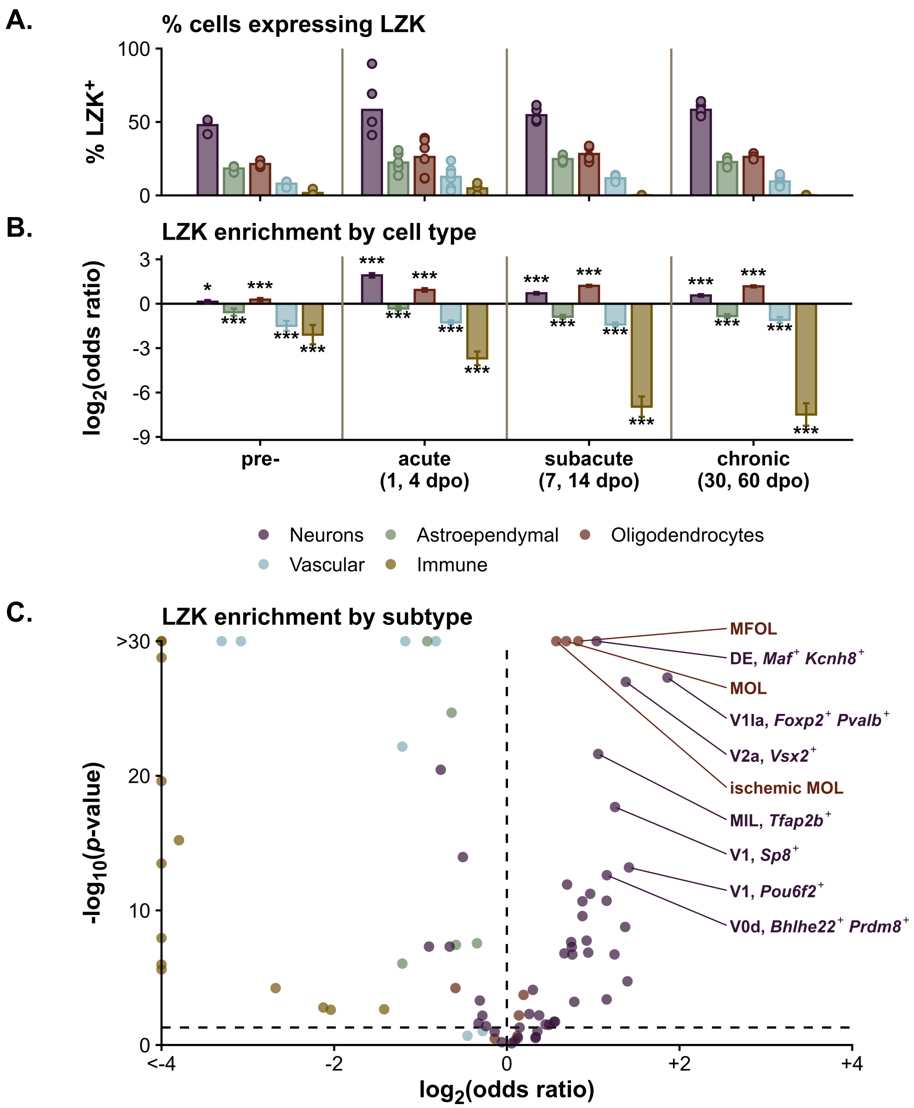


**Figure S3: LZK expression patterns in spinal cord cell types after SCI. (A)** Percentage of LZK-expressing cells (*Map3k13* > 0) across major cell types and SCI phases, derived from published single-nucleus RNA-Seq data. Each point represents the average per biological replicate; n = 2-3 female mice per time. Timepoints are grouped into uninjured, acute (1–4 days post-injury), subacute (7–14 days), and chronic (30–60 days). **(B)** Cell type–specific enrichment of LZK expression, estimated using a binomial generalized linear model. Plotted are $\log_2$ odds ratios (OR) with 95% confidence intervals. Asterisks indicate statistically significant enrichment compared to all other cell types (*$p < 0.05$, **$p < 0.01$, ***$p < 0.001$, ****$p < 0.0001$). **(C)** Subtype-level enrichment of LZK expression using the same modeling approach, aggregated across timepoints. Each point represents a distinct cell subtype; x-axis shows $-\log_{10}$(FDR-adjusted p-value), y-axis shows $\log_2$ odds ratio. The ten most significantly enriched subtypes (positive $\log_2$ OR) are labeled: ischemic MOL (ischemic mature oligodendrocytes), MFOL (myelin forming oligodendrocytes), MOL (mature oligodendrocytes), DE (dorsal excitatory), V1Ia/V1 (ventral inhibitory interneurons), V2a/VEP (ventral excitatory projection neurons), V0d (ventral commissural interneurons), MIL (medial excitatory, lateral). Marker genes noted where relevant.

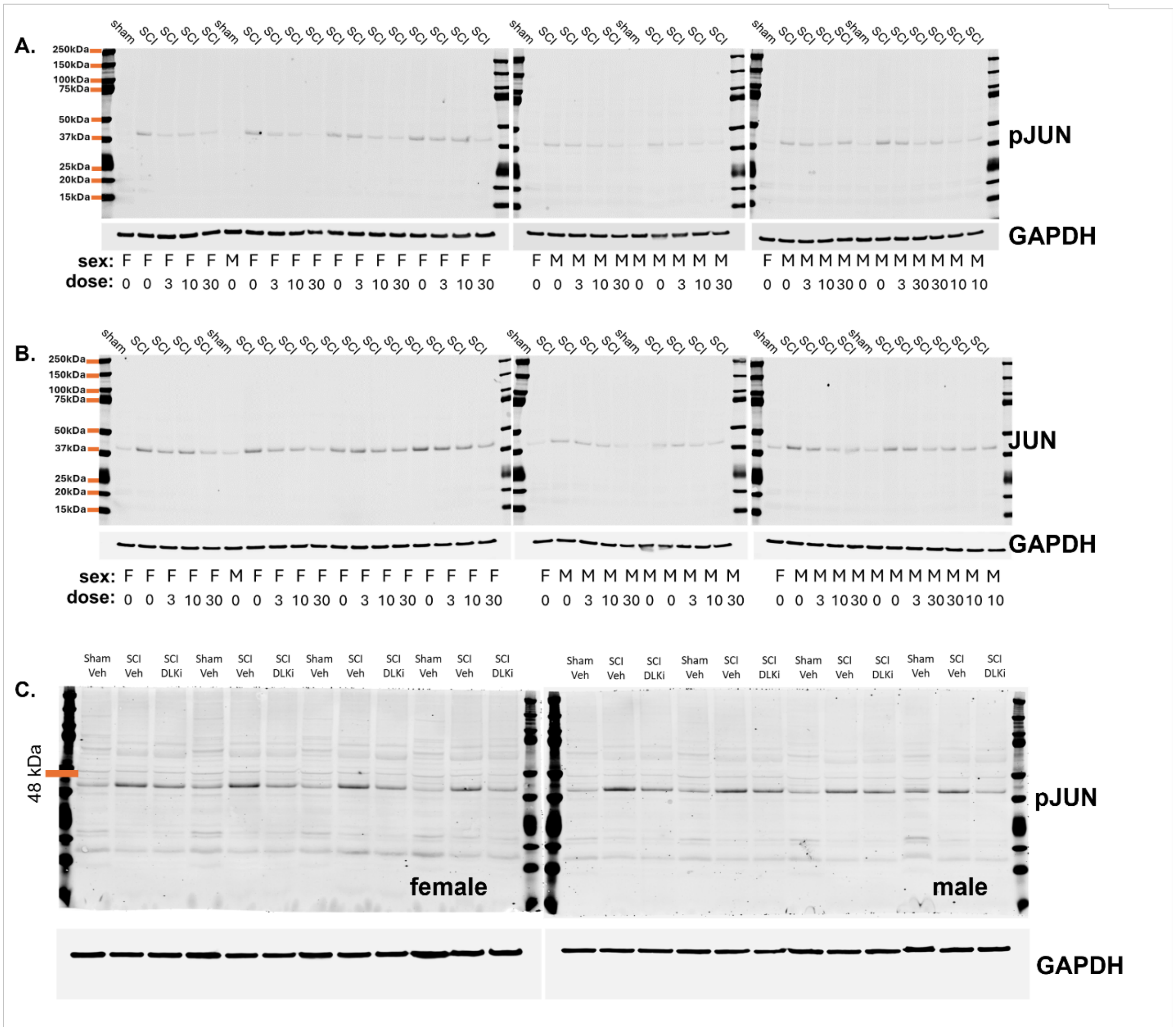


**Figure S4: Western blot images showing IACS-52825 suppression of JUN phosphorylation after SCI. (A, B)** Representative immunoblots for phospho-JUN (**A**) and total JUN (**B**) from spinal cord lesion epicenter tissue collected 24 hours after SCI and oral administration of IACS-52825 (3, 10, or 30 mg/kg) or vehicle. GAPDH was used as a loading control for each blot. Molecular weight markers are indicated in kilodaltons (kDa). Above each blot are annotations for treatment group (vehicle or IACS-52825 dose); below each lane are annotations for sex (F = female, M = male) and dose (in mg/kg). Quantification of normalized and scaled band intensities is shown in **Fig. 4A–B**. (**C**) Representative immunoblots for phospho-JUN from spinal cord lesion epicenter tissue collected 48 hours after SCI and a single intraperitoneal (i.p.) injection of vehicle or IACS-52825 (30 mg/kg). GAPDH was used as a loading control. Annotations above each blot indicate surgery type and treatment group. Lanes on the left correspond to female samples; lanes on the right correspond to male samples. Quantification of normalized and scaled band intensities is shown in **Fig. 4D**.

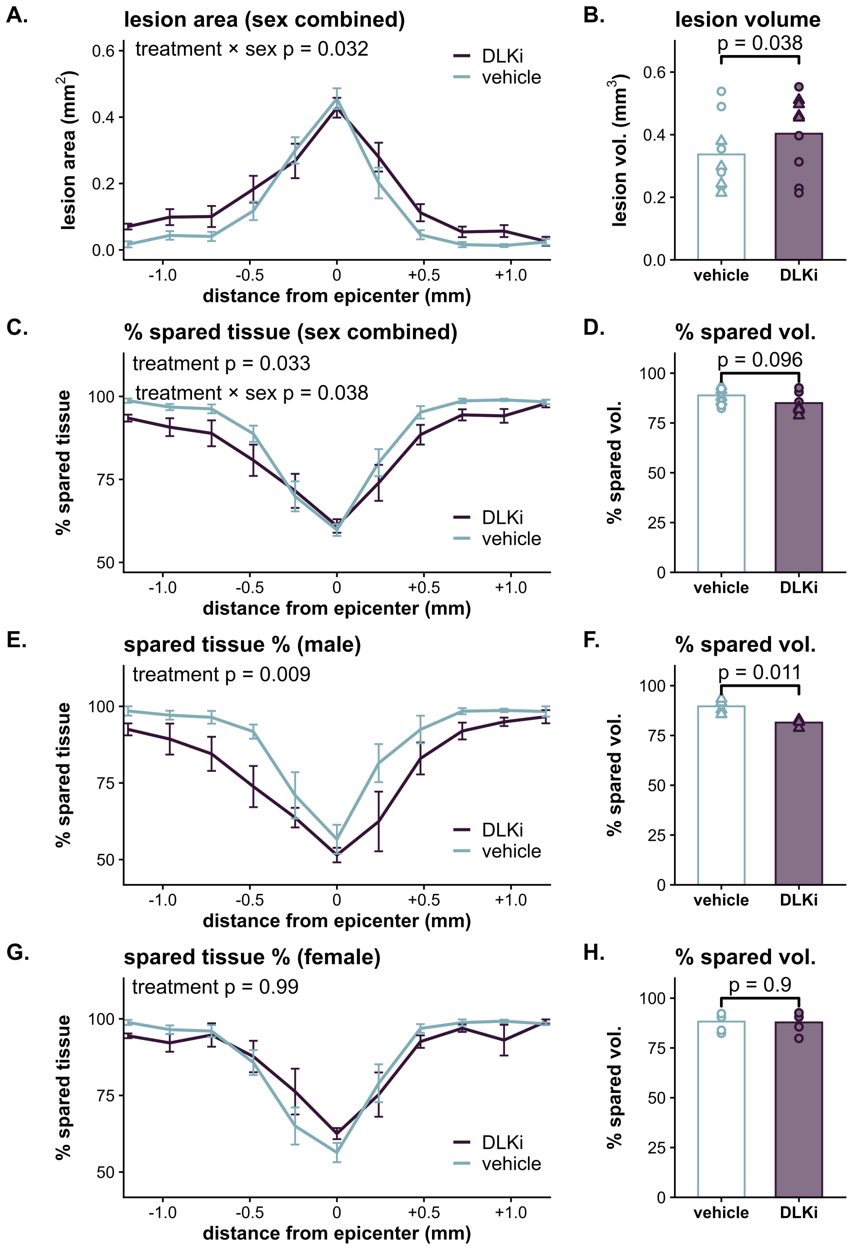


**Figure S5: Lesion area and spared tissue quantification after IACS-52825 treatment. (A)** Lesion area across serial spinal cord sections ±1.2 mm from the epicenter, averaged across sexes. A treatment × sex interaction was detected by linear mixed-effects modeling (p = 0.032). **(B)** Lesion volume calculated from area measurements in **A. (C)** Percent spared tissue, calculated as 100 × (total section area – lesion area) / total section area, across the same serial sections. Significant main effect of treatment (p = 0.033) and treatment × sex interaction (p = 0.038) detected by linear mixed-effects modeling. **(D)** Spared tissue volume calculated from percent spared tissue in **C. (E)** Percent spared tissue across serial spinal cord sections in males. A significant main effect of treatment was detected by linear mixed-effects modeling (p = 0.009). **(F)** Spared tissue volume in males; compared by unpaired t-test. **(G)** Percent spared tissue in females; no significant treatment effect detected (p = 0.99). **(H)** Spared tissue volume in females; compared by unpaired t-test. Triangles indicate males, circles indicate females. Error bars represent SEM. n = 5 females and 4 males per group.

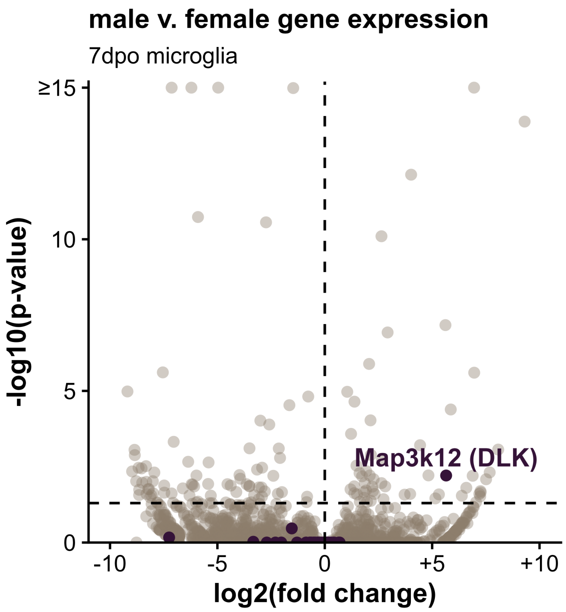


**Figure S6: Differential gene expression between male and female microglia at 7 dpo**. Volcano plot showing differential expression results from pseudobulked microglia at 7 days post-injury, comparing male and female mice in the *Tabulae Paralytica* dataset (GSE234774). Counts were aggregated per replicate using Seurat's *AggregateExpression* function, and differential expression was tested using DESeq2 with a Wald test. Each point represents a single gene; the x-axis shows $\log_2$ fold change (male vs. female), and the y-axis shows $-\log_{10}$(FDR-adjusted p-value). Global multiple testing correction was applied across all layers and cell types. DLK pathway genes and known DLK targets are labeled in purple. *Map3k12* (DLK) is significantly upregulated in male microglia ($\log_2$FC = 5.7, FDR-adjusted p = 0.006).
